# Quantifying Molecular Bias in DNA Data Storage

**DOI:** 10.1101/566554

**Authors:** Yuan-Jyue Chen, Christopher N. Takahashi, Lee Organick, Kendall Stewart, Siena Dumas Ang, Patrick Weiss, Bill Peck, Georg Seelig, Luis Ceze, Karin Strauss

## Abstract

DNA has recently emerged as an attractive medium for future digital data storage because of its extremely high information density and potential longevity. Recent work has shown promising results in developing proof-of-principle prototype systems. However, very uneven (biased) sequencing coverage distributions have been reported, which indicates inefficiencies in the storage process and points to optimization opportunities. These deviations from the average coverage in oligonucleotide copy distribution result in sequence drop-out and make error-free data retrieval from DNA more challenging. The uneven copy distribution was believed to stem from the underlying molecular processes, but the interplay between these molecular processes and the copy number distribution has been poorly understood until now. In this paper, we use millions of unique sequences from a DNA-based digital data archival system to study the oligonucleotide copy unevenness problem and show that two important sources of bias are the synthesis process and the Polymerase Chain Reaction (PCR) process. By mapping the sequencing coverage of a large complex oligonucleotide pool back to its spatial distribution on the synthesis chip, we find that significant bias comes from array-based oligonucleotide synthesis. We also find that PCR stochasticity is another main driver of oligonucleotide copy variation. Based on these findings, we develop a statistical model for each molecular process as well as the overall process and compare the predicted bias with our experimental data. We further use our model to explore the trade-offs between synthesis bias, storage physical density and sequencing redundancy, providing insights for engineering efficient, robust DNA data storage systems.

---

Storing data in DNA is attractive due to its information density of petabytes of data per gram, and excellent durability^1^. High-throughput (HT) sequencing and synthesis technologies^2,3^ have evolved and made storing information in synthetic DNA an increasingly realistic alternative to traditional long-term storage methods ^4–7^. However, the sequencing coverage of oligonucleotide (henceforth referred to simply as “oligo”) copy distribution was found to be very uneven, requiring modern error correction codes capable of handling sequence dropout^1,6–11^. Current methods typically require either trial-and-error on experimental protocols or brute-force use of hundreds to thousands sequencing reads to capture underrepresented sequences. This inefficiency stems from lack of understanding about bias in oligo copy distribution, as well as how it changes as the oligos are manipulated in DNA data storage systems.

The problem with quantifying bias in a DNA storage system is that we cannot immediately and clearly distinguish bias created by DNA synthesis from bias caused by PCR and sequencing. As our first foray in separating bias effects stemming from DNA synthesis versus PCR, we tagged an arbitrarily chosen DNA archival file with over 400,000 sequences using Unique Molecular Identifiers (UMI), random barcodes to label each molecule^12^. UMI labeling allowed us to decouple synthesis bias from PCR bias, and we found significant bias from DNA synthesis. To corroborate this finding, we ordered from Twist Bioscience a carefully designed ready-to-sequence pool with 1,536,168 sequences, each of which unique, and already containing necessary segments of DNA to be sequenced. This ready-to-sequence pool can be sequenced using an Illumina sequencer directly, with no need for intermediate PCR or DNA ligation steps required for sequencing library preparation. Thus we can quantify synthesis oligo distribution without any interference from molecular processes. To the best of our knowledge, this is the first time an oligo pool from array-based synthesis is characterized in this way. We found that synthesis bias was highly related to spatial location of oligos on a synthesis chip.

After quantifying synthesis bias, we studied PCR bias from two sources – guanine/cytosine (GC) content and PCR stochasticity. GC content of individual sequences had been previously found to affect PCR amplification efficiency in biological DNA^13–15^. In DNA storage, the GC content of each strand is determined by a data-to-DNA sequence encoder. We tested GC bias using two different oligo pools: one pool was encoded to avoid all homopolymers (non-homopolymer pool); in contrast, the other was encoded without homopolymer avoidance steps (homopolymer pool). These two encoding strategies indeed led to different GC distributions: the GC content of the homopolymer pool was between 25% and 75%, while that of the non-homopolymer pool was between 40% and 60%. However, in either case, we found no practically important association between GC content and PCR bias. Instead, we found that PCR stochasticity widened oligo copy distributions of our test DNA archival file and, based on our observations, seemed to be a dominant factor in PCR bias. PCR is an exponential process, so small random variations early on in amplification can have a large impact on distribution^16–22^.

Based on these observations, we constructed a computational model for predicting molecular bias in a DNA data storage system (Fig. 1). We have observed strong association between the bias predicted from this model and from our experimental data. Furthermore, we used our model to investigate the tradeoffs between synthesis bias, physical redundancy for storing DNA (i.e., oligo copy number) and sequencing redundancy (i.e., sequencing coverage). A system model can be very useful to determine the best parameters for a given DNA storage system.

**Figure 1.**
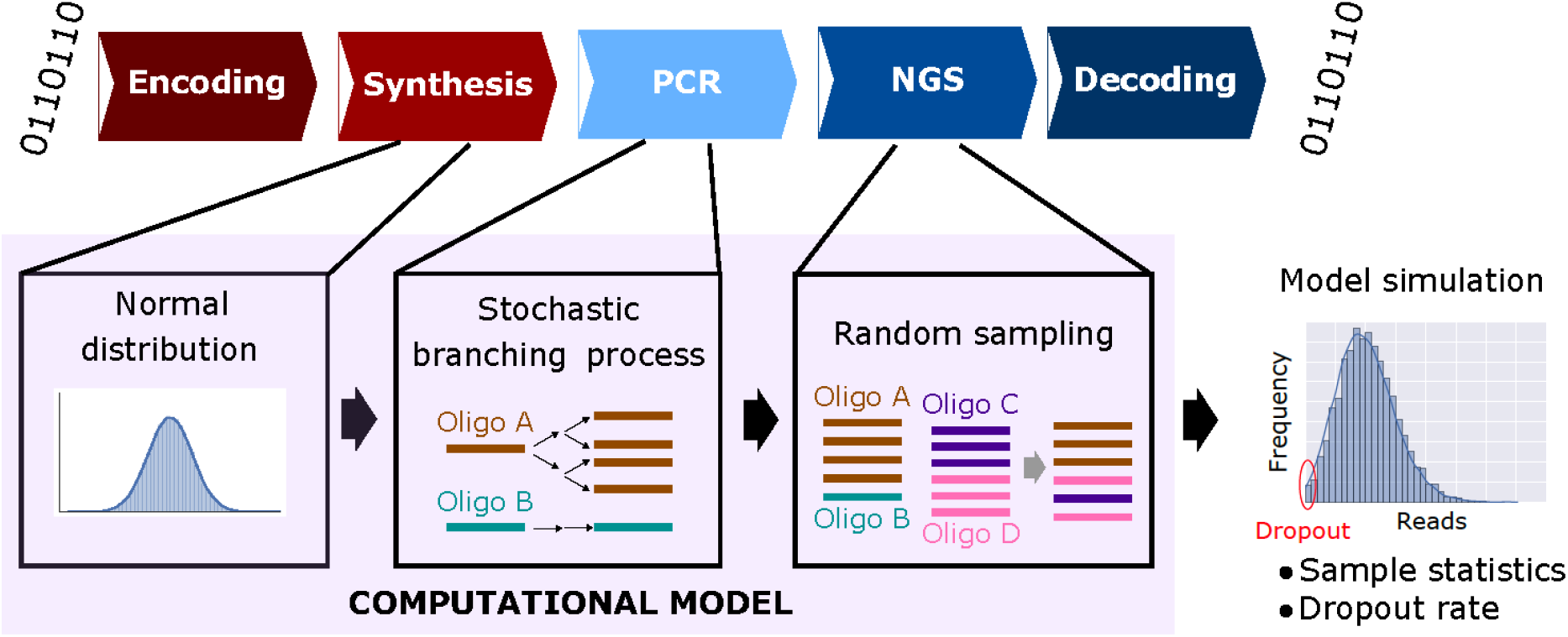
A DNA storage system model. Digital information is first encoded into oligonucleotide (oligo) sequences, resulting in multiple 150-base DNA fragments synthesized using array-based DNA synthesis technology, which are then stored. To read back the stored data, target DNA oligos can be selectively (random-) accessed using polymer chain reaction (PCR), then sequenced via next generation sequencing (NGS), and decoded back to digital information. The computational model approximates each molecular process in the DNA storage system: it uses a normal distribution for modeling sequence copy numbers from synthesis, a stochastic branching process for PCR, and random sampling for sequencing. The computational model makes predictions for oligo copy distribution to help researchers estimate statistics such as sequence dropout rate.

## UMI reveals that DNA synthesis is a prominent source of sequence bias in DNA data storage

Determining the source of bias in DNA data storage, and more generally in arbitrary DNA pools, is complicated because synthesis bias and PCR bias are typically coupled. To decouple them, we applied Unique Molecular Identifiers (UMI), barcodes to individually identify each molecule of an initial pool, in our case an arbitrarily chosen DNA file with over 400,000 sequences (Fig. 2a and **Supplementary Fig. S1**). Synthetic DNA pools include multiple copies of each sequence, and UMI labeling ensures with high probability that each molecule will include a tag different from any other. The UMI-labeled oligos were sequenced, and the resulting reads were aligned to the file sequences in two manners. First, these reads were aligned to individual sequences in the file using BWA^23^, independent from UMI, and their respective counts (coverage) are reported in Fig. 2c. Second, the same reads were aligned to sequences in the file, then further filtered by UMI label (Fig. 2b), and finally reported in Fig 2d. The UMI-filtered results are a proxy for the oligo distribution after DNA synthesis, and the copy number is clearly variable. This distribution is also very similar in shape to the distribution after PCR, indicating that PCR does not significantly increase bias overall. Nevertheless, PCR still has an impact on individual sequence counts, so we decided to study the *amplification ratio* of each sequence as a function of the number of initial molecules representing it. We define the amplification ratio to be the ratio of total reads after PCR to UMI count (i.e., oligo count before PCR) for each sequence. Fig. 2e shows that regardless of the initial oligo copy number, the *average* amplification ratio remains constant. On the other hand, the amplification ratio was observed to have high variation when oligos had very low copy numbers, indicating that the amplification ratio was affected by stochastic effects at these low copy numbers. Indeed, since a PCR process is composed of successive rounds of binomially distributed copying (each molecule has some probability of being copied), we would expect the standard deviation (s.d.) of the amplification ratio to be inversely proportional to the square root of the initial number of strands. Additionally, since observation takes a sequencing reaction (another binomial process) we’d expect a constant amount of added deviation. These observations lead us to the model:

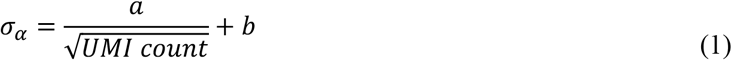

where *σ_α_* is the s.d. of the amplification ratio, and *a* and *b* are constants. Our experimental data was fitted using equation 1 and shown in Fig. 2f.

**Figure 2.**
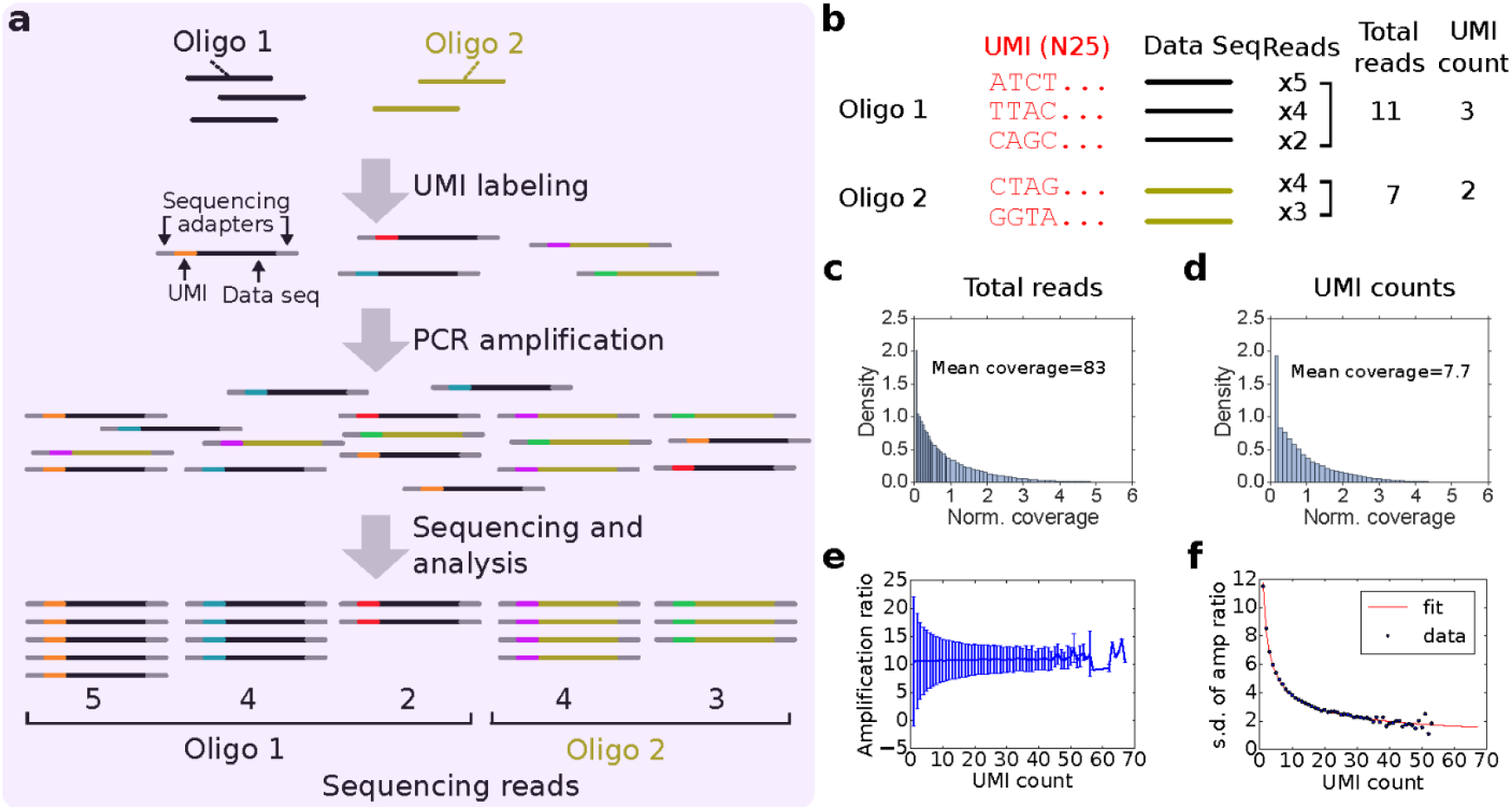
Estimating oligonucleotide bias using unique molecule identifiers (UMIs). (**a**) Overview of tagging each single DNA molecule with UMIs. Each oligo sequence (e.g., represented in black, beige) in a pool has multiple copies and each copy is labeled with a UMI (represented in different colors) and universal Illumina sequencing adapters (represented in grey). After UMI labeling, oligos are PCR-amplified and sequenced. (**b**) Hypothetical examples of UMI counting. The UMI count of each sequence is a proxy for the oligo copy number from DNA synthesis. The total number of reads containing the same UMI is a proxy for the number of copies of a DNA molecule created by PCR. (**c**) The distribution of number of reads for each sequence, normalized to 83.0 mean coverage. Read counts are normalized to form a probability density (y-axis); the integral of the probability density is 1 (see Methods). (**d**) The distribution of UMI counts for each sequence, normalized to 7.7 mean coverage. The biased UMI count distribution indicates that pools are already biased immediately after DNA synthesis, before any PCR is performed. (**e**) Amplification ratio versus UMI count. The average amplification ratio is roughly constant across UMI counts, but oligos with low initial copy numbers show higher variation. (**f**) Standard deviation (s.d.) of amplification ratio versus UMI count. The experimental data agree with equation 1.

## Synthesis bias is related to the spatial location on the synthesis chip

To further understand the synthesis bias, we ordered a carefully designed a *ready-to-sequence pool* with 1,536,168 unique DNA sequences from Twist Bioscience. Oligos in this pool already contain universal Illumina adapters and Illumina sequencing primers on both 5’ and 3’ ends, allowing us to sequence it without any sequencing library preparation such as PCR or ligation. By mapping the sequencing reads of each sequence back to its corresponding location on the synthesis chip, a distinct pattern can be observed (Fig. 3a, left), indicating that synthesis bias was related to the spatial location on the synthesis chip. After further discussion with Twist Bioscience, their synthesis process was improved, and the oligo counts on the synthesis chip became much more even (Fig. 3b, left). Interestingly, the oligo distribution before the synthesis process improvement did not follow a normal distribution, but the oligo distribution using the improved synthesis process is now well fitted to a normal distribution (Figs. 3a and 3b, right).

**Figure 3.**
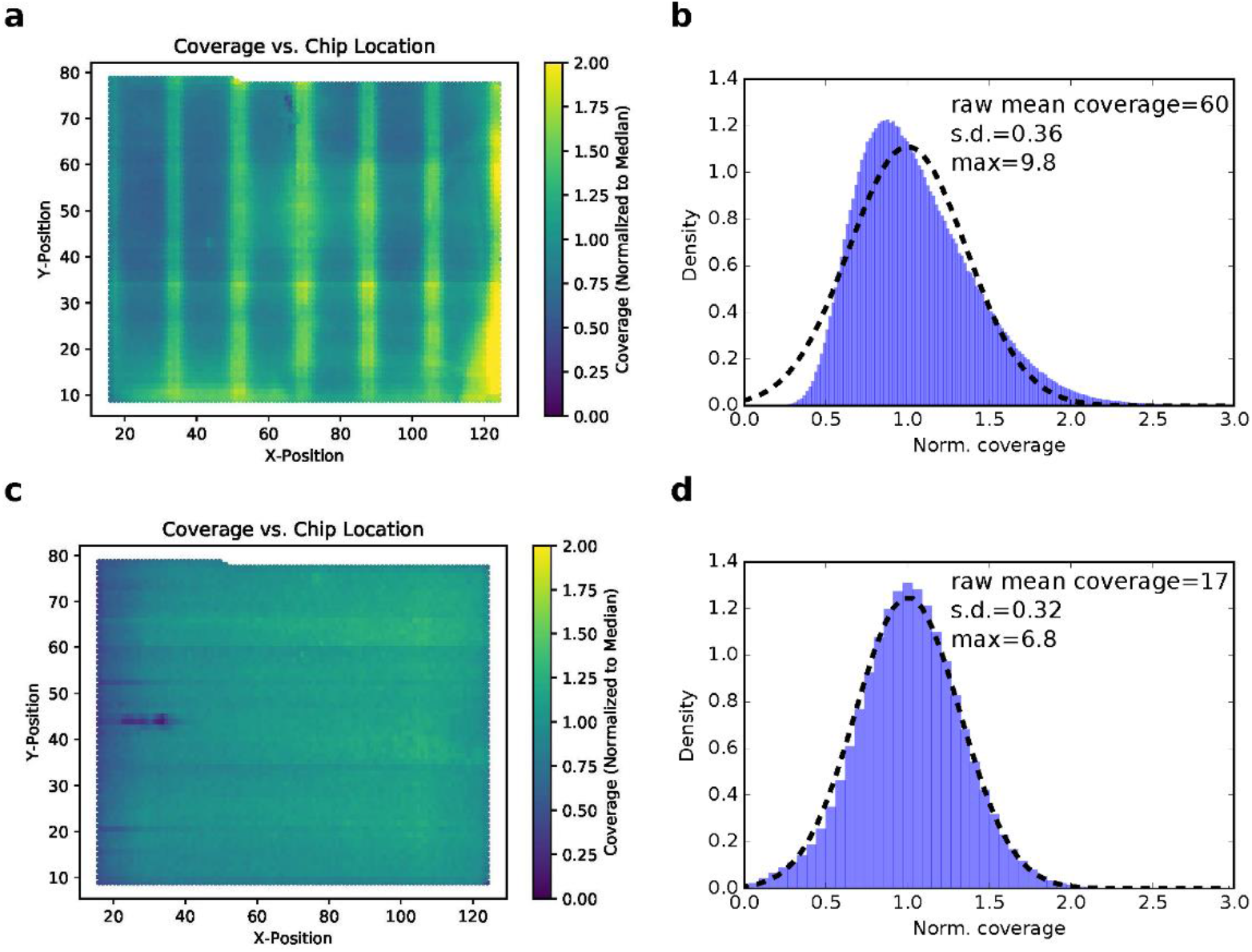
Oligo copy distribution on the synthesis chip. (**a**) The sequencing coverage of each oligo from the first ready-to-sequence pool was mapped back to its corresponding location on the synthesis chip. Coverages are normalized to the median. (**b**) The histogram of normalized sequencing coverage of the first ready-to-sequence pool (blue). The distribution does not fit a normal distribution (dashed line). (**c**) The sequencing coverage of each oligo from the second ready-to-sequence pool, mapped back to its corresponding location on the synthesis chip. Coverages are normalized to the median. (**d**) The histogram of normalized sequencing coverage of the second ready-to-sequence pool (blue). The second ready-to-sequence pool showed much more even oligo copy distribution and fits a normal distribution (dashed line) much more closely than then first.

## Population fraction change for quantifying PCR bias

We now turn to studying the PCR bias by creating metrics to quantify it at the sequence level. We begin by defining the *population fraction* of a sequence *i* after *k* ∈ [*Z*_≥0_] cycles of PCR as

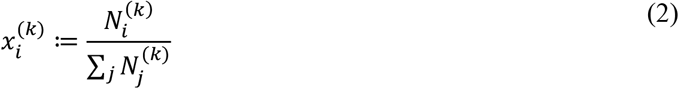

where 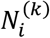 is the number of reads of sequence *i* after *k* PCR cycles. We then define the *population fraction change* for sequence *i* to be

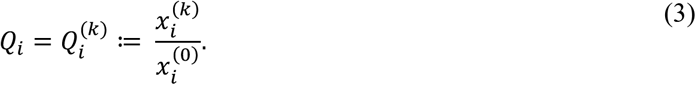

We consider a PCR process to be *unbiased* when 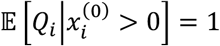 for all sequences, that is, no sequence becomes over or underrepresented after an unbiased PCR, whereas we consider a PCR process to be *biased* when 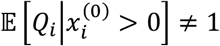 for any sequence *i*. We then can say that experiments with higher standard deviation over the population fraction change, s. d. 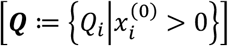, show more bias when all other conditions are equivalent. It is worth noting that even an unbiased process will have s. d. [***Q***] > 0 for finite sample sizes. Furthermore, s. d. [**Q**] should asymptotically decrease with the total number of reads.

## PCR bias is not correlated with GC content in DNA data storage

Although previous studies observed PCR bias in genomic biological sample amplification^13–15^, it remains unclear whether such bias is significant in DNA data storage. To assess this, we used the 1.5 million-sequence ready-to-sequence pool and compared its distribution before PCR and after PCR. The ready-to-sequence pool was sequenced in two ways: (1) directly from the synthesized pool and (2) after one 6-cycle plus five 5-cycle PCR processes, for a total of 31 cycles. Each PCR process was limited to no more than 6 cycles to prevent resource exhaustion (i.e., there was always an excess of primer and other reagents). Sequencing data (Fig. 4a) shows qualitatively little change in the coefficient of variation (c.v.) of oligo copy distribution before and after PCR (0.41 and 0.45, respectively, when both are subsampled to 20x coverage). The two datasets were then compared at a sequence level by observing *population fraction changes* with respect to the overall available the pre-PCR pool coverage, 60x (Fig 4b). The distribution before PCR shows the effect of subsampling on population fraction, and the distribution after PCR shows the effect of PCR itself. The latter showed much higher standard deviation. The standard deviations of *population fraction changes* were 0.24 and 0.37 before PCR and after PCR, respectively, and these two numbers were statistically different (p < 0.005, computed by bootstrapping n=1000). This indicates that PCR increased bias relative to a random sampling process.

**Figure 4.**
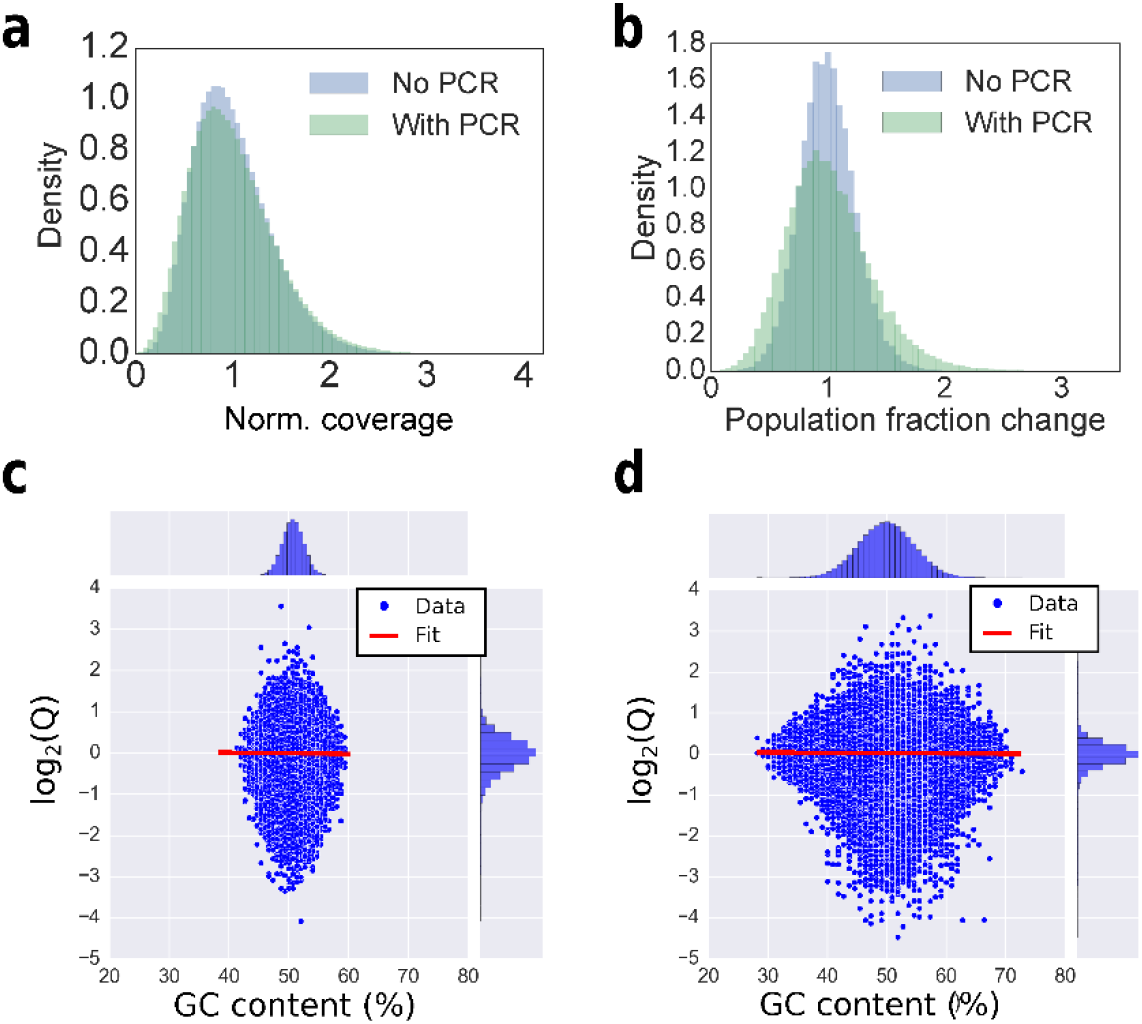
Impact of GC content on PCR bias. (**a**) Histogram of sequencing coverages for optimized ready-to-sequence pool. The ready-to-sequence pool was sequenced directly, without PCR (blue). The same pool was amplified using PCR for 31 cycles (green) and sequenced separately. Both were randomly sampled to coverage of 20x for direct comparison, and they look quite similar (c.v. = 0.41 and c.v.=0.45, respectively). (**b**) The blue histogram shows the *population fraction change Q* distribution of the ready-to-sequence pool, before PCR, after being subsampled to 20x coverage, with respect to its overall available coverage (60x). The green histogram shows the *population fraction change Q* distribution of the ready-to-sequence pool, after PCR and after being subsampled to 20x coverage, with respect to the pre-PCR pool distribution at 60x coverage. The blue distribution shows the effect of subsampling, while the further widening of the green curve with respect to the blue curve is attributed to stochastic bias in the PCR process. (**c, d**) The GC content is plotted against the log2 of the *population fraction change Q* for (**c**) the ready-to-sequence, non-homopolymer pool and a homopolymer pool (**d**). The experimental data are shown as blue dots, and the linear fit is shown as a red line. The histograms of GC content and *log_2_*(*Q*) are shown at the top and right, respectively.

We then asked whether *population fraction changes* were caused by GC content. We first examined the ready-to-sequence pool, which was encoded to avoid homopolymers^6^ (Fig. 4c). Although the association between *population fraction changes* and GC content of this pool (between 40% and 60%) was found to be statistically significant (P value < 0.05), the association between the two was very small and practically unimportant (the slope of the linear fit was < 0.01). Additionally, we tested another 9 different DNA archival files with a total of 1,358,998 unique sequences that allow random homopolymers (Fig 4d; **Supplementary Figure S2** shows experimental workflow details). These homopolymer files had a wider range of GC content from 25% to 75%, but the association between GC content and the *population fraction changes* was still very small and not practically important (the slope of the linear fit was < 0.01). The negligible bias impact from GC content in our experimental data was likely because these oligos were relatively short (150-nt), and the use of KAPA HIFI polymerase also reduced the impact of GC bias^24^. Having established that GC content was not the main effect being observed, we turned to hypothesizing that PCR stochasticity was the culprit.

## PCR stochasticity can lead to significant bias

Because PCR is not perfect (i.e., replication of an individual molecule has a probability of less than one), even small random divergence in early phases of amplification can create significant bias, which is known as PCR stochastic bias. We have shown that PCR bias is related to oligo copy number in the UMI quantification experiment, especially for sequences with low copy numbers in the initial pool (from a previous PCR process or from a biased synthesis pool). Now we want to understand better how PCR stochastic bias affects our DNA storage system.

To quantify PCR stochastic bias, we used an arbitrarily chosen DNA pool with 7,373 sequences to perform a serial dilution-PCR experiment (Fig. 5a). The master pool was diluted to different average copy numbers ranging from 8 to 113 (the copy numbers were quantified using qPCR). Then each sample was amplified with 18 cycles of PCR using primers with Illumina sequencing primer overhangs. Subsequently, a second step of PCR was carried out to include the Illumina adapters where we adjusted the number of cycles to equalize the final library concentration (**Supplementary Figure S3** shows workflow details). The second PCR was carried out at high copy number of the templates (over a million oligo copies per sequence) to avoid introducing additional bias. Our experimental results show that as average copy number decreased, oligo distribution skewed further away from its mean (Fig. 5c). We plot average copy number in a pre-PCR mix against the coefficient of variation (c.v.) of sequencing coverage (Fig. 5d) and standard deviation of *population fraction change Q* (**Supplementary Figure S4**). Both plots show that the lower oligo copy numbers were, the greater the PCR stochastic bias was.

**Figure 5.**
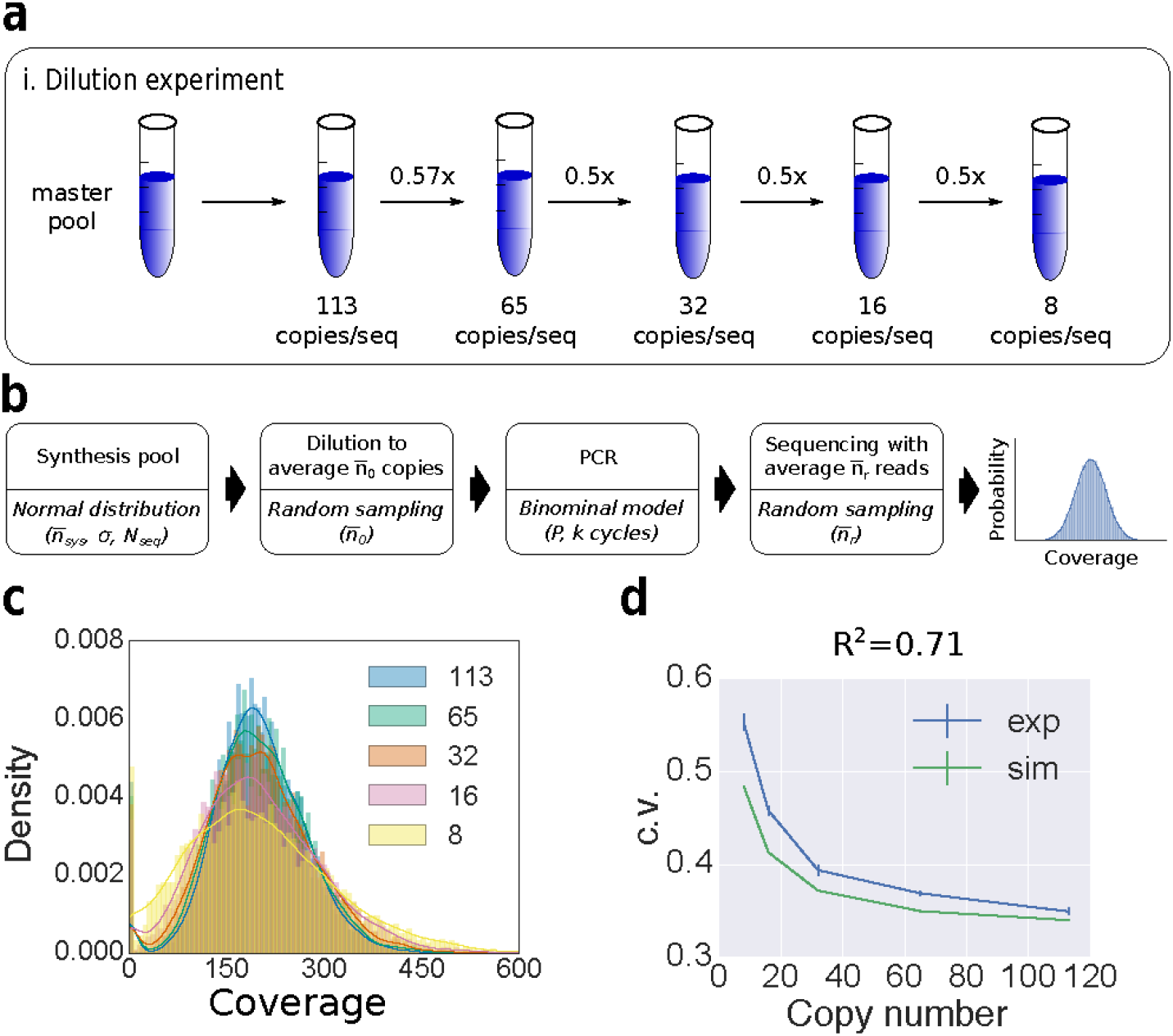
Dilution-PCR experiment. (**a**) The experimental workflow. A master DNA pool was diluted to different average copy numbers as indicated in the drawing. Each dilution sample was PCR-amplified and sequenced using an Illumina NextSeq instrument, and the results sampled at 200x coverage. (**b**) A computational model for the dilution-PCR experiments. The synthesis pool model used *N_Seq_* = 7,373 number of sequences, and normally distributed copy numbers with mean 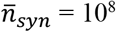, and standard deviation *σ* = 3.2 * 10^7^. The c.v. of the synthesis pool in this simulation 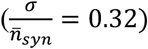 was set to be equal to the c.v. of our ready-to-sequence pool sequenced at mean coverage 17. The dilution process was simulated using random sampling with a mean copy number 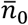, ranging from 8 to 113. PCR was simulated as a binomial process with a probability of successful amplification *P* = 0.95 and 18 PCR cycles. The simulated sequencing result was obtained using random sampling with an average coverage 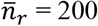. (**c**) Simulated post-PCR sequencing coverage histogram of each dilution-PCR sample. The initial (pre-PCR) average copy number of each histogram is shown in the legend, ranging from 8 to 113. Coverage counts are normalized to display a probability density. A Gaussian estimated density curve is added as an outline of each histogram to help with visualization. (**d**) Sequencing coverage c.v. of the post-PCR mix versus average copy number in the pre-PCR mix. The model prediction (green) shows good agreement with the experimental data (blue) with R^2^=0.71. The error bars of experimental data indicate standard error calculated from triplicate experiments. The error bars of model outputs indicate standard error calculated from 100 repeated simulations.

## A computational model can predict molecular bias in a DNA archival system

After characterizing the bias caused by synthesis and PCR sequencing retrieval, we construct a DNA storage model that encompasses the entire workflow of DNA storage, starting from synthesis ➔ aliquot into pre-PCR reaction ➔ PCR amplification with *k* cycles ➔ sequencing with mean 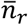 reads (Fig. 5b). We model the oligo copy distribution of synthesis as a normal distribution with total number of sequences *N_seq_*, mean copy number per sequence 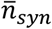, and standard deviation of oligo copy number *σ*. The PCR process is modeled as a stochastic branching process using the following recursive equation:

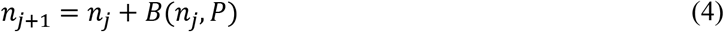

where *n_j_* is the number of molecules in the *j-th* cycle; *B(n_j_, P)* is a binomially distributed random variable with *n_j_* molecules, and *P* is the probability of a successful amplification. Illumina sequencing was previously observed to have bias on GC extreme sequences^13,25,26^, but GC content in our files did not show practically significant bias in the PCR GC bias test. Therefore, high-throughput sequencing and sample dilution are modeled using random sampling. Note that for performance reasons our model does not perform stochastic simulation for high copy number PCR because PCR carried out at high copy number of templates should obey the law of mass action and therefore be effectively deterministic.

We then interrogated our computational model to determine whether it can estimate the bias observed in the serial dilution-PCR experiment. Despite not being able to observe the oligo population directly after synthesis, our UMI experiment (Fig. 2) has provided evidence that its population distribution is quite similar to the distribution resulting from a PCR process that starts from a large average copy count sample coming from that synthesized pool. As such, the copy distribution of a synthesis pool is modeled as a normal distribution with the same c.v. as the experimental data from the (optimized) ready-to-sequence pool. Then we used our system model to simulate the dilution-PCR experiment. Fig. 5d shows that our model prediction is in good agreement with the c.v. of the experimental data (R^2^ = 0.71). The model also predicted the trend of standard deviation of *population fraction change Q*: the lower starting copy number in the PCR showed higher standard deviation (R^2^ = 0.84; **Supplementary Figure S4**).

## A computational model can help determine system parameters for DNA data storage systems

Taking it one step further, we used our computational model to study a range of parameters associated with DNA storage: synthesis bias, physical redundancy for storing DNA, and sequencing redundancy (Fig. 6a). In particular we investigated the impact of these parameters on sequence dropout rate, which is critical for error-free decoding. Fig. 6b plots sequence dropout rates as a function of the c.v. of a synthesis pool and sequencing reads. It shows that a biased synthesis pool (i.e., high c.v.) is the dominant factor in sequence dropout and cannot be proportionally compensated by additional sequencing reads. Sequence dropout is caused by physical storage with a limited number of oligo copies coupled with PCR stochastic bias. Fig. 6c plots sequence dropout rates as a function of the copy number of stored DNA and sequencing reads. It shows that physical storage density is a more important factor than sequencing reads in modulating sequence dropout. Interestingly, our model estimates that it is possible to store as few as 10 copies per oligo sequence (information density of 9.3 EB/g (EB: exabytes; 10^18^ bytes)), while achieving less than 2% sequence dropout. This estimated information density is over 10-fold higher than prior work by Erlich and Zielinski^9^, showing that there is more optimization to be done in this area.

**Figure 6.**
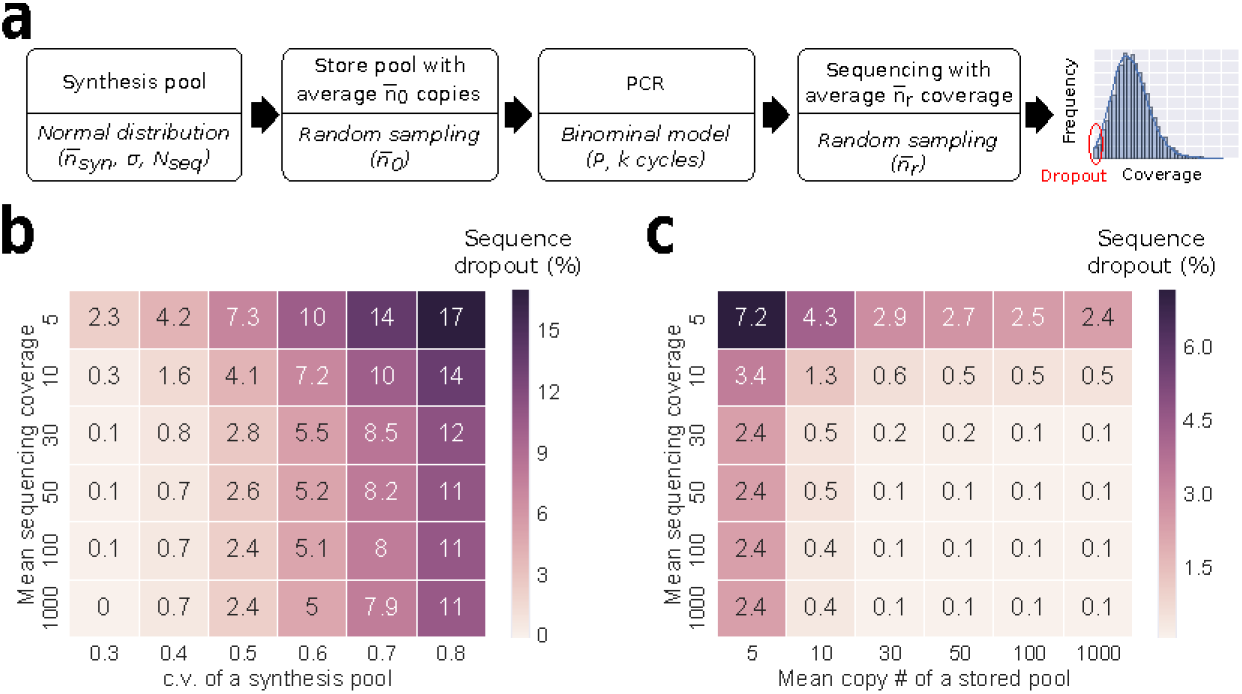
A computational model can help determine system parameters for DNA data storage. (**a**) A synthesis pool was generated with *N_seq_* = 10,000 total number of sequences, with normally distributed copy numbers with a mean of 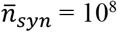 and standard deviation *σ* = 3.2 * 10^7^. The pool was simulated to store an average copy number 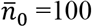, followed by 20 cycles of PCR amplification with *P*=0.95, and high throughput sequencing with average sequencing coverage 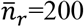. Sequence dropout (i.e., coverage of 0 for a given sequence) rates were quantified. (**b**) Sequence dropout percentage as a function of variable synthesis pool c.v. and variable mean sequencing coverage 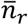. (**c**) Sequence dropout percentage as a function of variable mean copy number *n_0_* and variable mean sequencing coverage 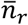. The reported dropout percentage was the average of 100 repeated simulations.

## Discussion

In this work, we quantified molecular bias in a DNA storage system, and we identified two significant bias sources: synthesis bias and PCR stochastic bias. Synthesis bias was found to be related to the spatial location on the synthesis chip, and this observation was later used to inform and improve the synthesis process. PCR stochastic bias was identified as the second main driver of oligo copy variation. Indeed, prior work also found that PCR copy data from a deeply diluted oligo pool resulted in dramatic bias, which was not suitable for data recovery^9^.

Another important contribution of this manuscript is the construction of the first process-wide model that provides a quantitative understanding of how oligo copy distribution is skewed as it goes through a DNA storage system. Importantly, such system model helps researchers rationally optimize the use of DNA storage density and sequencing redundancy for reliable data decoding without conducting hundreds of experimental trials. We believe this is an important step towards engineering robust, efficient DNA storage systems.

Our system model was experimentally tested by PCR-amplifying a single file without any other non-targeted files in a pool. This experiment was designed to avoid complexity from other files for proper quantification of the impact of PCR stochastic bias. Next, we plan to investigate whether PCR random access of a file from a complex pool with additional files will lead to more bias. We suspect that amplifying a very small file from a complex pool with relatively large number of sequences will exhibit more copy number variation due to non-specific binding of primers. New methods will probably be needed for such system.

## METHODS

### Reagents

All DNA pools were synthesized by Twist Bioscience (San Francisco, CA). All DNA pools were resuspended to 10 ng/μL in 1X TE buffer (pH 7.5). All primers were purchased as desalted, unpurified DNA from Integrated DNA Technologies (IDT; Coralville, IA). All primers were resuspended to 100 uM in 1X TE buffer (pH 7.5). KAPA HIFI polymerase was purchased from Kapa Biosystems. T4 ligase and T4 Polynucleotide Kinase (T4 PNK) were purchased from New England Lab.

### PCR protocol

In a 20 μL PCR reaction, 1 μL of 1 ng/μL of ssDNA pool was mixed 1 μL of 10 uM of the forward primer and 1 μL of 10 uM of the reverse primer, 10 μL of 2X KAPA HIFI enzyme mix, and 7 μL of molecular biograde water. The reaction followed a thermal protocol: (1) 95°C for 3 min, (2) 98°C for 20 sec, (3) 62°C for 20 sec, (4) 72°C for 15 sec. After PCR, the length of the PCR products was confirmed using a Qiaxcel fragment analyzer, and the sample concentration was measured using a Qubit 3.0 fluorometer.

### Sample preparation for sequencing

Before sequencing, the concentrations of all samples were quantified using qPCR. The final sample was then prepared for sequencing by following the NextSeq System Denature and Dilute Libraries Guide. The final concentration of the loaded sample for our Illumina NextSeq is 1.3pM, and a 10%-20% PhiX was spiked in as a control (PhiX is a genomic DNA sample provided by Illumina).

### Protocols of UMI labeling

The general workflow for UMI labeling of a single-stranded DNA pool is divided into 5 steps (**Supplementary Figure S1**): (1) phosphorylation of a ssDNA pool and Illumina P7 adapters, (2) assembly of a ssDNA pool with Illumina adapters with DNA staples by heat annealing, (3) ligation of Illumina adapters to the ssDNA pool, (4) extraction of the ligated sample using denaturing polyacrylamide gel electrophoresis (D-PAGE), and finally (5) PCR enrichment of the full length product.

The phosphorylation of ssDNA was performed using the following recipe: 5 pmole of the single-stranded DNA pool, 20 units of T4 Polynucleotide Kinase (T4 PNK), 1μL of 10X T4 ligase buffer and 1 μL of 10X T4 PNK buffer were mixed in a 10 μL total volume reaction. 500 pmole of single-stranded Illumina P7 adapter, 200 units of T4 PNK, 5μL of 10X T4 ligase buffer and 5μL of 10X T4 PNK buffer were mixed in a 50 μL total volume reaction. The mixtures were incubated for 30 minutes at 37°C.

The assembly of the single-stranded DNA pool with adapters were performed with the following recipe: In a 25 μL reaction, 15 pmole of single-stranded DNA pool, 30 pmole of DNA staples and 45 pmole of Illumina P5 and P7 adapters were mixed. The mixture was heated up to 95 °C for 2 minutes, and then cooled down to 25 °C at a rate of 1 degree per minute.

Ligation of DNA was performed with a 15 μL reaction in which 10 μL of the assembled DNA mixture, 2 μL of the T4 ligase (10 units/μL), 1.5 μL of T4 ligase buffer and 1.5 μL of molecular water were mixed. The ligation mixture was incubated at room temperature for 30 minutes, followed by heat inactivation at 65°C for 10 minutes.

A 10% D-PAGE gel was made by mixing 2.5 mL of 19:1 40% acrylamide/bus, 1.2 mL of 10X TBE, 5.04 g of urea and deionized water to 12 mL. Then 72 μL APS and 4.8 μL of TEMED were added to help polymerization. DNA sample was mixed with 2X TBE/Urea denaturing loading buffer (Bio-Rad). Gels were run at 200 V for 55 minutes at 55°C. The extracted band was incubated with 1X TE buffer overnight at room temperature for elution.

The eluted single-stranded DNA was PCR-amplified using the end primers of Illumina adapters. The PCR reaction used 1 μL of the eluted single-stranded DNA, 10 pmole of the forward and reverse primers, 10 μL of 2X KAPA HIFI polymerase and 8 μL of molecular water. The thermal protocol is as follows: (1) 95°C for 3 min, (2) 98°C for 20 sec, (3) 60°C for 20 sec, (4) 72°C for 15 sec.

### Density histogram plots

The y-axis of a density histogram shows probability density, and the area (or integral) under the histogram is 1. The probability density *d_i_* is calculated by dividing the count by the sample size times its bin width (see the following equation).

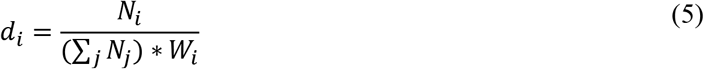

where *N_i_* is the count of the i-th bar, and *W_i_* is the bin width of the i-th bar. Displaying the y-axis as probability density makes it possible to compare distributions. In Fig. 5c, a Gaussian estimated curve is added to help visualize each histogram.

## ACKNOWLEDGEMENTS

We would like to thank Patrick Finn, Siyuan Chen, Andrew Stewart, Bernadette Arias, and Emily Leproust from Twist Bioscience for supplying us with the DNA and for suggestions on the analysis of DNA synthesis data. We also thank Leila Zelnick for discussion and comments on the manuscript.

## AUTHOR CONTRIBUTIONS

Y-J.C. designed, performed, and analyzed experiments. C.T. and Ke.S. designed experiments and analyzed data. L.O. performed experiments. S.D.A. ran the DNA sequence encoder and decoder, and analyzed data. G.S., K.S., and L.C. directed and supervised the work.

