## supplemental for "Quantifying Molecular Bias in DNA Data Storage"

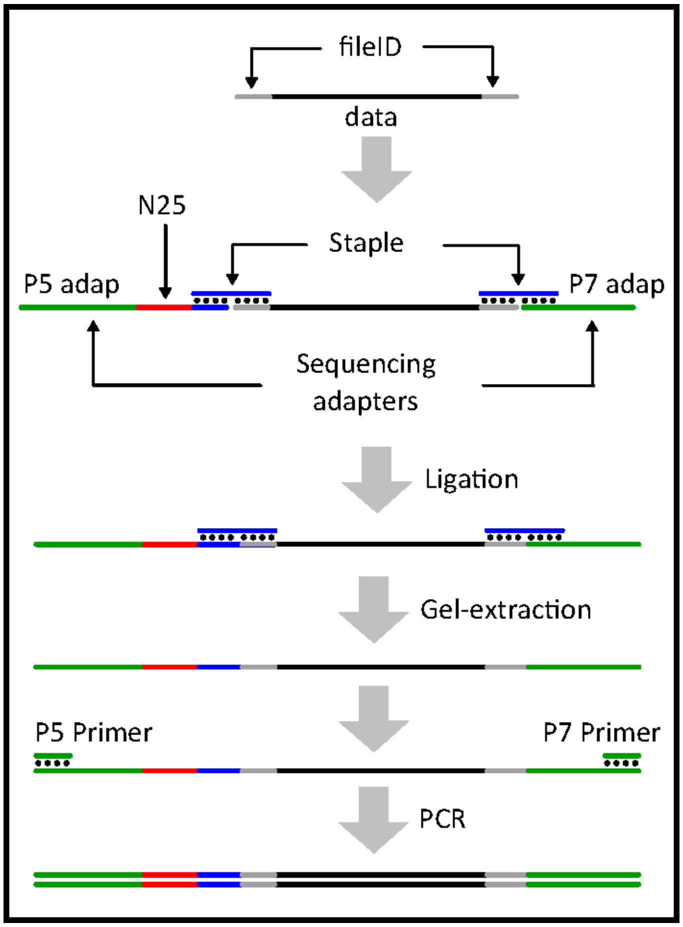


**Supplementary Figure S1.** DNA data strands are assembled with sequencing adapters using two staples. Note that one of the adapters contains a randomized region (N25) which serves as UMI. After assembly, DNA nicks are sealed using T4 DNA ligase. A D-PAGE gel is then used to extract the ligated strands. Finally, two end primers are used to enrich the full-length product for sequencing on an Illumina instrument.

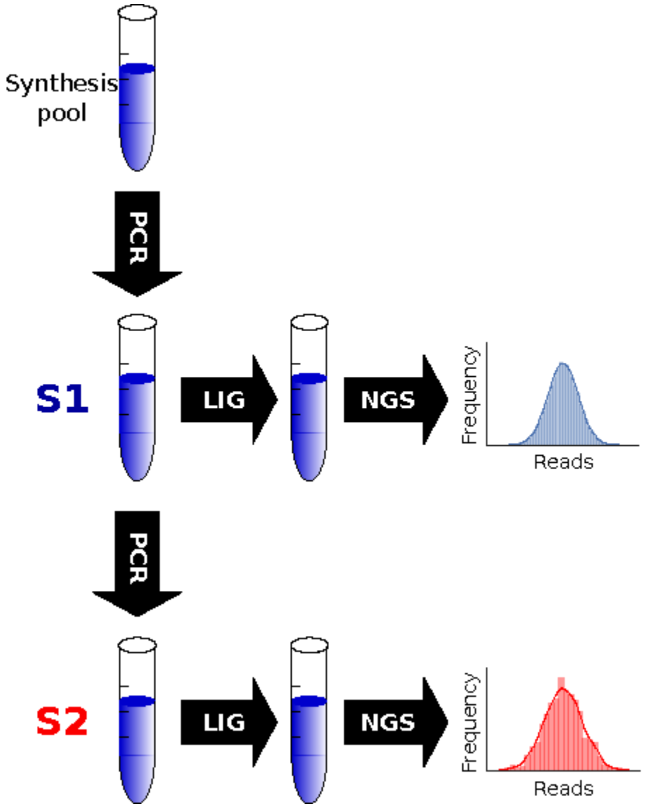


**Supplementary Figure S2.** To decouple PCR bias from synthesis bias, we used a 2-step PCR-sequencing method. The synthesis pool was PCR-amplified with a pair of primers to obtain an amplified file *S1*. *S1* is PCR-amplified again with the same primers to obtain *S2*. *S1* and *S2* were ligated to Illumina sequencing adapters and sequenced to get their oligo copy distributions. By comparing the distributions of *S1* and *S2*, we can quantify PCR bias.

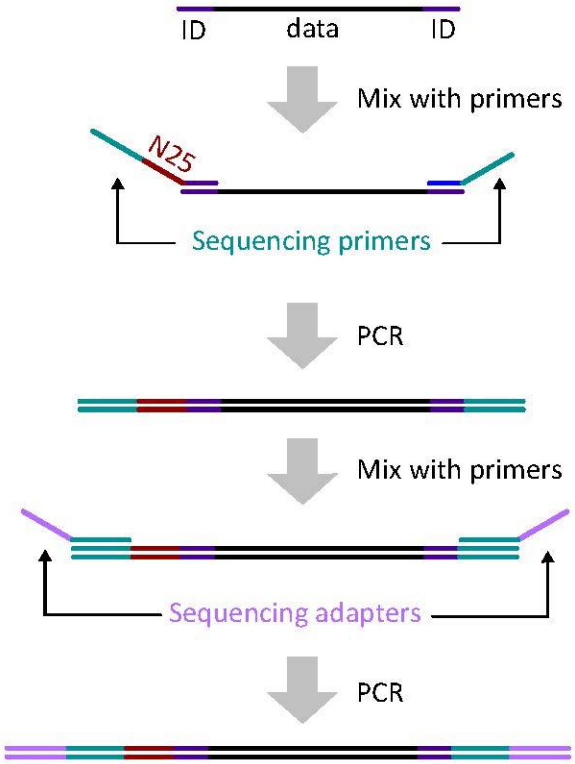


**Supplementary Figure S3.** Workflow of dilution-PCR experiments. A DNA pool is first amplified with primers that include Illumina sequencing primers overhangs and a randomized region (N25). The randomized region is used for increasing the diversity for an Illumina NextSeq instrument. After that, the amplified oligos were amplified with another set of primers with Illumina sequencing adapters.

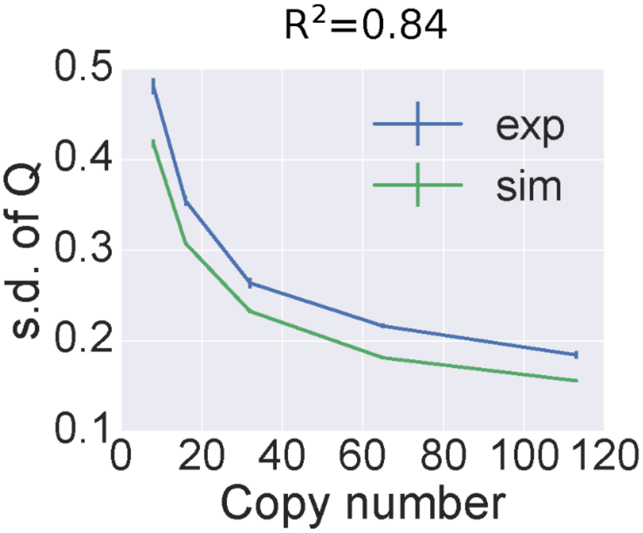


**Supplementary Figure S4.** Standard deviation of *population fraction change Q* of the post-PCR DNA versus average copy number of the pre-PCR mix. The blue trace shows the standard deviation of Q in the post-PCR experimental data sequenced and then sampled at 200x coverage where the pre-PCR mix contains an average of 8 to 113 copies per sequence and its Q is calculated comparing to the same mix at average 200 copies per sequence. The green trace shows the simulated data, i.e., the standard deviation of Q in the post-PCR mix after simulated sequencing and sampling at 200x coverage where the pre-PCR mix contains an average of 8 to 113 copies per sequence and its Q is calculated comparing to the same mix at average 200 copies per sequence with c.v. = 0.32. The model prediction (green) captures a trend similar to the experimental data (blue) with $R^2$=0.84. The error bars of the experimental data indicate standard error calculated from triplicate experiments. The simulation is plotted in green color, and the errors indicate standard error from 100 repeated simulations.

**Python code for simulating a DNA storage system (Fig. 6).**

```
%matplotlib inline
import matplotlib.pyplot as plt
import seaborn as sns
from collections import defaultdict
import pysam
import numpy as np
import seaborn as sns
import pandas as pd
import seaborn as sns

def bootstrapSamp(counts_runA, cov_tot):
    np.random.seed()
    pb_runA = counts_runA/np.sum(counts_runA)
    _q =
np.random.choice(range(0,len(counts_runA)),size=cov_tot,replace=True,p=pb_run
A.astype(np.float))
    sim = np.zeros(len(pb_runA))
    for idx in _q:
        sim[idx] += 1
    return sim

def compute_FFC(base_run, pcr_run, include_nan=False):
    a = base_run/np.sum(base_run)
    b = pcr_run/np.sum(pcr_run)
    old_settings = np.seterr(divide='ignore',invalid='ignore')
    ampratio = b/a
    if include_nan == False:
        ffc = ampratio[np.isfinite(ampratio)]
    else:
        ffc = ampratio
    _=np.seterr(**old_settings)  # reset to default
    return ffc

## PCR sim
def binormSim(n, p, cyc=10):
    # simulate the number of molecules after "c" cycle
    # n: number of template molecules
    # p: probability of success PCR
    for i in range(cyc):
        n += np.random.binomial(n, p, 1)
    return n.item(0)

# simulate CV of synthesized oligos vs sequencing coverage
nseqs = pow(10,4)
p = 0.95
nsyn = pow(10,8)
dropout_list = []
store_copyN = 100
for SEQCov_mean in [5,10,30,50,100,1000]:
  for CV in [0.3, 0.4, 0.5, 0.6, 0.7, 0.8]:
      sigma = store_copyN*CV
      s_normal = np.random.normal(nsyn, sigma, nseqs)
      syn_pool = np.array([round(c) if c>0 else 1 for c in s_normal])
      store_pool = bootstrapSamp(syn_pool, store_copyN*nseqs)
```

```
      simPCR_counts1 = np.array([binormSim(int(count), p, 20) for count in
store_pool]).astype(float)
      bsSEQ_counts = bootstrapSamp(simPCR_counts1, SEQCov_mean*nseqs)
      missing_counts = np.count_nonzero(bsSEQ_counts==0)
      dropoutRate = 100.0*missing_counts/nseqs
      dropout_list.append((SEQCov_mean, CV, dropoutRate))

df_dpRate = pd.DataFrame(np.array(dropout_list), columns = ["Sequencing
reads", "Oligo pool CV", "dropoutRate"])
df_dpRate["dropoutRate"] = df_dpRate["dropoutRate"].astype(float)
df_dpRate["dropoutRate"] = df_dpRate["dropoutRate"].apply(lambda x:
round(x,1))
df_dpRate["Sequencing reads"] = df_dpRate["Sequencing reads"].astype(int)
df_dpRate["Oligo pool CV"] = df_dpRate["Oligo pool CV"].astype(float)

df_dpRate_T = df_dpRate.pivot("Sequencing reads", "Oligo pool
CV","dropoutRate")
sns.set_context("notebook", font_scale=2.0, rc={"lines.linewidth": 2.5})
f, ax = plt.subplots(figsize=(9, 6))
sns.heatmap(df_dpRate_T, annot=True, linewidths=.5, ax=ax)


# simulate storage copy number vs sequencing coverage
nseqs = pow(10,4)
nsyn = pow(10,8)
CV = 0.32
p = 0.95
dropout_list = []
for store_copyN in [5, 10, 30, 50, 100, 1000]:
  for SEQCov_mean in [5, 10, 30, 50, 100, 1000]:
      sigma = nsyn*CV
      s_normal = np.random.normal(nsyn, sigma, nseqs)
      syn_pool = np.array([round(c) if c>0 else 1 for c in s_normal])
      store_pool = bootstrapSamp(syn_pool, store_copyN*nseqs)
      simPCR_counts1 = np.array([binormSim(int(count), p, 20) for count in
store_pool]).astype(float)
      bsSEQ_counts = bootstrapSamp(simPCR_counts1, SEQCov_mean*nseqs)
      missing_counts = np.count_nonzero(bsSEQ_counts==0)
      dropoutRate = 100.0*missing_counts/nseqs
      dropout_list.append((store_copyN, SEQCov_mean, dropoutRate))

df_dpRate = pd.DataFrame(np.array(dropout_list), columns = ["Copy # per seq",
"Sequencing reads", "dropoutRate"])
df_dpRate["dropoutRate"] = df_dpRate["dropoutRate"].astype(float)
df_dpRate["dropoutRate"] = df_dpRate["dropoutRate"].apply(lambda x:
round(x,1))
df_dpRate["Copy # per seq"] = df_dpRate["Copy # per seq"].astype(int)
df_dpRate["Sequencing reads"] = df_dpRate["Sequencing reads"].astype(int)
df_dpRate_T = df_dpRate.pivot("Sequencing reads", "Copy # per seq",
"dropoutRate")
sns.set_context("notebook", font_scale=2.0, rc={"lines.linewidth": 2.5})
f, ax = plt.subplots(figsize=(9, 6))
sns.heatmap(df_dpRate_T, annot=True, linewidths=.5, ax=ax)
```

| Name | Sequences | Length (bp) |
|---|---|---|
| P5 adapter | AATGATACGGCGACCACCGAGATCTACACTCTTTCCCTACACGACGCTCTTCCGATCTNNNNNNNNNNNNNNNNNNNNNNNNNAGTGAGGTAGAGGTGTATTC | 103 |
| P7 adapter | GATCGGAAGAGCACACGTCTGAACTCCAGTCACATCACGATCTCGTATGCCGTCTTCTGCTTG | 63 |
| P5 staple | TGCTGGTAACAACTTGCCTTGAATACACCTCTACCTCACT/3SpC3/ | 40 |
| P7 staple | AGACGTGTGCTCTTCCGATCTTGGTTTGATTACGGTCGCA/3SpC3/ | 40 |
| P5 primer | AATGATACGGCGACCACCGAGA | 22 |
| P7 primer | CAAGCAGAAGACGGCATACGAG | 22 |

**Supplementary Table 1. Sequences of UMI labeling.** /3SpC3/ represents a C3 spacer modification at the 3'end.

| Pool | File | Forward Primer | Reverse Primer | File Size (bytes) | Number of sequences |
|---|---|---|---|---|---|
| Ready-to-sequence pool | | AATGATACGGCGACCACCGAGA | CAAGCAGAAGACGGCATACGAG | 10,686,220 | 1,536,168 |
| Homopolymer Pool | 1 | TCCTGCTTGCGTTAAATGGA | TTCCGCAAGACTTATTGGCA | 5,043,000 | 241,648 |
| Homopolymer Pool | 2 | ACCGCGCTCGAAGAATTTAA | TCGCAACACCTTTCGTACAA | 6,295,900 | 301,680 |
| Homopolymer Pool | 3 | AAACAAAGTTAGCGGCTCGT | AGGCCGCGAATTTGGATTAT | 5,815,000 | 278,640 |
| Homopolymer Pool | 4 | AATTTGGCATTCACCGTGGA | AGTCGCCAAATAAGTGCCAT | 6,690,000 | 320,565 |
| Homopolymer Pool | 5 | AGCCTTGTGTCCATCAATCC | ATTAGCCAAACCATAGCGCA | 600,200 | 28,761 |
| Homopolymer Pool | 6 | TGTATTTCCTTCGGTGCTCC | AAACCAGACCGTTGTCGAAA | 573,700 | 27,491 |
| Homopolymer Pool | 7 | TGTGTTCCTCCTCGGTATGA | AGGAAGCGCCAACTAATTGT | 180,500 | 8,650 |
| Homopolymer Pool | 8 | TAGCCTCCAGAATGAAACGG | TACACACGGTTTGGCTTGAA | 3,163,000 | 151,563 |
| Serial dilution PCR experiment (Fig. 5) | | ACATTCCGTGCCATTGGATT | TTTGTGGAACGATTTGCCGA | 115,394 | 7,373 |

**Supplementary Table 2. Pools/files used in this study.** All encoded pools/files are listed. All files were encoded in 150-base DNA strands.
